# VirTarget: Virus-Informed Pharmacogenomics Framework Identifies Immunotherapies That Mitigate Epstein–Barr Virus–Driven Dysregulation in Multiple Sclerosis

**DOI:** 10.1101/2025.11.03.686445

**Authors:** Bekir Altas, Polymnia Georgiou, Marios Onisiforou, Panos Zanos, Anna Onisiforou

**Affiliations:** Department of Biomedical Sciences, Marquette University, Milwaukee, WI, United States; Department of Psychology, University of Wisconsin Milwaukee, Milwaukee, WI, United States; S.K. Lifestyle Technologies LTD, Nicosia, 2048, Cyprus; Department of Psychology, University of Cyprus, Nicosia, 2109, Cyprus; Center of Applied Neuroscience, Nicosia, 1678, Cyprus

**Keywords:** Multiple sclerosis, Epstein–Barr virus, Virus-informed pharmacogenomics, Immunotherapies

## Abstract

**Background:** Epstein–Barr virus (EBV) is increasingly recognized as a central driver of multiple sclerosis (MS), yet current immunotherapies are selected without regard to their effects on EBV or EBV–MS genetic interactions. To address this gap, we developed *VirTarget*, the first *virus-informed pharmacogenomics network framework* that systematically evaluates approved MS immunotherapies with respect to EBV-driven pathogenesis and host genetic susceptibility risk.

**Methods:** *VirTarget* integrates three complementary layers: (1) EBV–host interactomics, mapping viral–host protein– protein interactions within MS-relevant pathways; (2) host genetic susceptibility, linking MS-associated variants to EBV-targeted, therapy-modulated networks; and (3) transcriptomics directionality analysis, contrasting drug signatures with the convergent MS–EBV transcriptomic signature to classify therapies as reinforcers or reversers.

**Results:** Network analysis revealed substantial heterogeneity among therapies. Dimethyl fumarate showed the strongest and broadest engagement with the MS–EBV network, followed by natalizumab and interferons, while anti-CD20 antibodies and S1P modulators exerted the weakest effects. Genetic mapping highlighted convergence on key risk genes (*HLA-DRB1, IL7R, IL2RA, CD40, TYR*) directly targeted by EBV proteins within therapy-modulated pathways. Transcriptomic profiling stratified therapies into reinforcers (e.g., dimethyl fumarate, fingolimod, dexamethasone, interferon-β1b) and reversers (e.g., prednisolone, cladribine, interferon-β1a, rituximab), reflecting opposing influences on the MS∩EBV signature. Notably, multiple therapies reversed EBV-driven dysregulation of cytokine and innate immune pathways, suggesting their benefits extend in counteracting viral-mediated modulation of immune processes.

**Conclusion:** *VirTarget* provides the first systems-level, virus-informed comparative map of how MS immunotherapies engage with the MS–EBV network, intersect with MS genetic susceptibility risk, and modulate both MS- and EBV-related transcriptomic signatures. By revealing drug-specific patterns of viral pathway engagement and genetic convergence, this framework establishes a foundation for precision treatment strategies in MS, where therapeutic selection is informed by viral serostatus, host genetics, and system-level transcriptomic responses.

## 1. Introduction

Multiple sclerosis (MS) is a chronic autoimmune and neurodegenerative disease affecting over 2.8 million people worldwide^1^. It is characterized by demyelination, neuroinflammation, and progressive loss of neurological function, and is the leading cause of neurological disability in adults aged 18–40 ^2^. The disease course is highly heterogeneous. Most patients initially present with clinically isolated syndrome (CIS), a first demyelinating event suggestive of MS, before progressing to relapsing–remitting MS (RRMS) and, over time, to secondary progressive MS (SPMS) ^3,4^. Reported conversion rate from CIS to definite MS ranges from 30% to 82%, depending on diagnostic criteria and follow-up duration ^3,4^. In contrast, a minority of patients (approximately 10–15%) develop primary progressive MS (PPMS), characterized by a gradual accumulation of disability from onset without distinct relapses or remissions ^5^. Despite decades of research, the precise etiology of MS has remained elusive, reflecting its complex interplay of genetic, environmental, and immunological risk factors.

Accumulating evidence strongly implicates Epstein–Barr Virus (EBV) as a necessary environmental risk factor for MS. EBV infects over 95% of adults globally, establishing lifelong latency within memory B cells ^6,7^. In a longitudinal study, EBV seroconversion conferred a >30-fold increased risk of developing MS, while nearly all patients with MS are EBV-seropositive ^8^. Importantly, the timing of EBV infection influences disease risk: primary infection in adolescence or adulthood, often manifesting as infectious mononucleosis, significantly increases MS susceptibility risk ^9^. This has shifted the view of MS from a purely autoimmune condition to a virus-driven disorder of immune dysregulation.

Genetic predisposition interacts closely with EBV biology. Genome-wide association studies have identified more than 200 MS risk loci, with the strongest signal arising from *HLA-DRB1\**15:01 ^10^. Carriers of this allele who mount antibodies against the EBV nuclear antigen 1 (EBNA1, residues 385–420) face a 24-fold increased risk of MS ^11,12^. Intriguingly, *HLA-DRB1\**15:01 has also been shown to function as a co-receptor for EBV entry into B cells ^13^, suggesting a direct mechanistic link between genetic susceptibility and viral persistence. Mechanistic studies further demonstrate that molecular mimicry between EBNA1 and CNS proteins such as GlialCAM and α-crystallin ^14,15^ can generate cross-reactive immune responses, allowing autoreactive T and B cells to escape tolerance and target CNS tissue. EBV-infected B cells may act as aberrant antigen-presenting cells, amplifying T cell activation and autoantibody production ^16^, while EBV persistence within meningeal ectopic lymphoid follicles provides chronic antigenic stimulation that fuels progressive forms of the disease ^17^. EBV also manipulates host immune signaling through viral–host protein–protein interactions (PPIs), such as modulating Th17 differentiation ^18^, a pathway that contributes to CNS inflammation and demyelination in MS ^19^.

These insights have reframed MS as a multifactorial disease in which EBV persistence, host genetics, and immune dysregulation act synergistically to trigger and sustain pathogenesis. This model also clarifies why MS relapses often occur following viral infections ^20–23^ and why disease activity differs between relapsing and progressive phases. In RRMS, systemic immune responses and environmental stressors may promote EBV reactivation and acute inflammatory relapses ^24–27^, whereas in SPMS and PPMS, inflammation becomes compartmentalized within the CNS, where EBV-positive B cells in meningeal follicles, oxidative stress, and chronic microglial activation drive ongoing neurodegeneration ^17,28^.

Paradoxically, current MS therapies were developed primarily as immunosuppressive or immunomodulatory agents, despite EBV being largely controlled by cytotoxic T cells. This creates the “immunosuppression paradox” in MS, whereby suppression of immune surveillance may inadvertently permit EBV reactivation or persistence. For instance, natalizumab prevents CNS immune infiltration but expands peripheral memory B cells, the major EBV reservoir, and has been linked to subclinical viral reactivation ^29,30^. Fingolimod impairs systemic immune surveillance and has been associated with EBV-positive CNS lymphomas ^31–33^. Alemtuzumab induces profound T cell depletion, impairing EBV-specific responses and predisposing to lymphoproliferative complications ^34–36^. In contrast, anti-CD20 therapies (rituximab, ocrelizumab) may exert their strong clinical efficacy by directly depleting EBV-infected memory B cells ^37–39^. Together, these findings highlight a critical gap, while EBV is implicated as a causal driver of MS, current treatments were designed primarily to suppress inflammation and immune activity rather than to target latent viral persistence within B cells. This underscores the need for therapeutic strategies that achieve both effective control of inflammation and restoration of antiviral immunity. There is therefore an urgent need to develop precision therapies that restore antiviral immunity against EBV or that directly disrupt EBV–host interactions underlying MS pathogenesis. Additionally, although inferences about the effects of MS immunotherapies can be made based on their mechanisms of action, we still lack a systematic understanding of how these therapies interact with the MS–EBV convergent signature, genetic susceptibility loci, and the viral–host pathways that sustain disease.

Previous studies have established the effectiveness of network-based methods to uncover viral-mediated pathogenic mechanisms in neurodegenerative and autoimmune diseases, including the role of EBV in MS ^18^, microbiota–virus–neurodegeneration interactions ^40^, the links between COVID-19 infection and neurological or neuropsychiatric disorders ^41^, and mechanistic connections between viral infections, including EBV, and Alzheimer’s disease (AD) ^42^. Network-based analyses has further proven effective in identifying shared molecular mechanisms and overlapping targetable pathways across comorbid conditions such as type II diabetes and neuropsychiatric disorders ^43^. Comparative analysis using network pharmacology have also evaluated the potential of existing antidiabetic medications to mitigate AD risk ^44^. Recently, we developed *VirTrack,* a computational framework that integrates EBV–host PPIs with MS clinical type–specific transcriptomes, moving beyond static network approaches to capture dynamic, clinical type-dependent viral influence across CIS, RRMS, SPMS, and PPMS ^45^. *VirTrack* revealed that EBV differentially targets immune, vascular, and lysosomal pathways across MS clinical types, underscoring the need for precision, stage-informed antiviral strategies ^45^.

Building on and extending this framework, we developed *VirTarget*, a virus-informed pharmacogenomic platform that integrates EBV–host interactomics, MS-associated proteins, genetic susceptibility, and transcriptomic signatures from MS, EBV, and drug responses. (**Figure 1**). This approach enables systematic, comparative analysis of all approved MS immunotherapies to determine how they engage EBV-modulated immune pathways, converge on genetic risk nodes, and rewire MS–EBV convergent transcriptomic signature, yielding the first comprehensive map of how current treatments reshape the molecular interface between EBV and MS pathogenesis. By uniting viral, genetic, and therapeutic dimensions, this study provides the first mechanistic and translational framework for resolving the immunosuppression paradox in MS and guiding the rational design of EBV-targeted precision immunotherapies.

**Figure 1:**
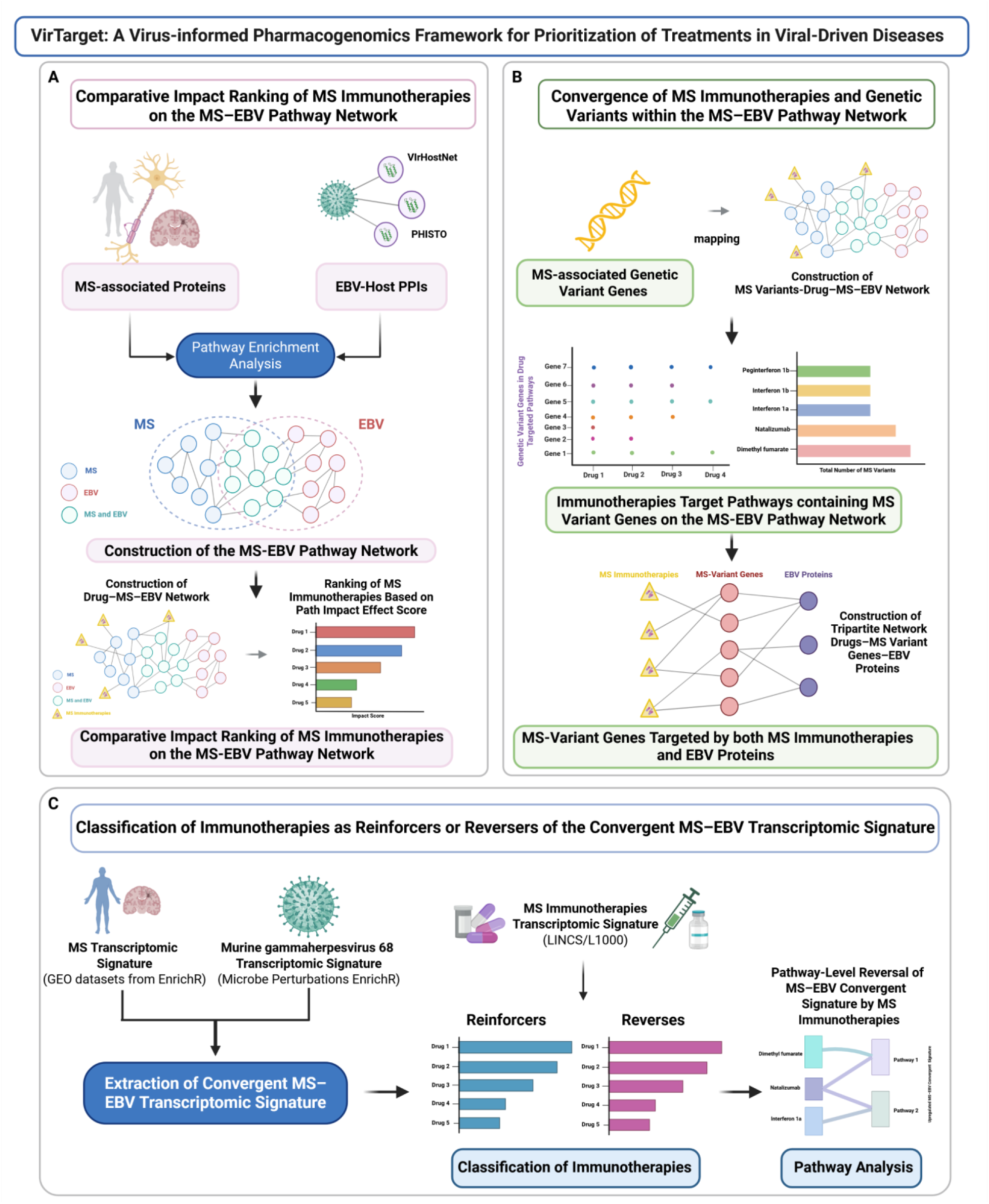
VirTarget: Virus-informed Pharmacogenomics Framework for Prioritization of Treatments in Virus-Driven Diseases. The VirTarget framework integrates three complementary layers of analysis to identify and prioritize immunotherapies that mitigate EBV–driven dysregulation in MS. **(A) Comparative impact ranking of MS immunotherapies on the MS–EBV pathway network.** MS-associated proteins and EBV–host PPIs were collected, followed by functional enrichment of MS and EBV datasets to construct the integrated MS–EBV pathway network. Drug–pathway interactions were then mapped, and a comparative impact analysis was performed by calculating the Disease Node Impact Effect Score, enabling quantification of each therapy’s effect within the network. **(B) Convergence of immunotherapies with MS genetic susceptibility.** MS-associated variant genes were mapped onto the MS–EBV pathway network to identify pathways containing MS risk variants that are also targeted by immunotherapies. A tripartite drug–gene–virus protein network was constructed to highlight MS-associated variant genes that are directly targeted both by immunotherapies (through drug–gene interactions) and by EBV proteins. **(C) Classification of immunotherapies as reinforcers or reversers of the convergent MS–EBV transcriptomic signature**. Drug-induced transcriptomic signatures (LINCS/L1000) were integrated with MS and EBV transcriptomic signatures to stratify therapies as either reinforcers (concordant) or reversers (discordant) of the MS–EBV convergent transcriptomic signature. Pathway-level enrichment analysis further identified pathways underlying reversal of the convergent MS–EBV transcriptomic signature by specific therapies. Abbreviations: EBV, Epstein–Barr virus; MS, multiple sclerosis; PPI, protein–protein interaction; LINCS, Library of Integrated Network-Based Cellular Signatures; L1000, LINCS gene-expression profiling platform.

## 2. Methods

### 2.1 Construction of the MS-EBV Pathway Network

We first reconstructed the MS-EBV pathway-pathway comorbidity network, following an established methodology from our previous works ^43,44^. This approach enables the identification of shared molecular mechanisms and potential therapeutic targets in comorbid diseases through the construction of disease-disease pathway-pathway networks. By using the *STRING disease* app in Cytoscape ^46^, we first collected the top 200 disease-associated proteins with the highest disease scores for MS (DOID:2377). We also collected experimentally validated virus-host PPIs for EBV from VirHostNet ^47^ and PHISTO ^48^ databases. Overall, 7073 virus-host PPIs were collected for three EBV strains (Human herpesvirus 4 strain B95-8 (taxid: 10377), EBV strain AG876 (taxid: 82830) and EBV strain GD1 (taxid 10376)) involving 1256 human targets and 153 viral proteins. Next, we performed enrichment analysis on the 200 disease-associated proteins of MS and the 1256 human proteins that are targeted by EBV using the Kyoto Encyclopedia of Genes and Genomes (KEGG) database. ClueGO ^49^ an app in Cytoscape, was used to conduct the enrichment analysis, considering only statistically significant pathways (adjusted *p*-value ≤ 0.05, Bonferroni step-down correction).

Using the significant KEGG pathways found for MS and EBV, we constructed the MS-EBV pathway-pathway network, where edge interactions between the pathways represent functional relationships between the pathways, that were collected form the KEGG database (Homo sapiens) using the KEGGREST package ^50^ in R. The final MS-EBV network consists of 98 pathways (nodes) and 328 edges, representing functional interactions between pathways. Among these, 15 pathways are shared between MS and EBV, highlighting common molecular mechanisms.

Based on a methodology we developed in our previous work ^44^, we also assigned weights to the edges of the constructed network to reflect their biological significance. Specifically, edges involving interactions between intersection nodes, which are pathways that can be found in both diseases, were assigned a weight of 3. Additionally, edges that involve interactions between an intersection and non-intersection nodes received a weight of 2. Edges that represent functional interactions between MS and EBV pathways were also assigned a weight of 2. All other edges were assigned a weight of 1. This weight assignment aimed to prioritize interactions between nodes from the two diseases that play a more significant role in facilitating the interaction between the two diseases.

The weighting scheme was designed to prioritize interactions with higher biological relevance. While no universal standard exists for assigning edge weights in disease-disease networks, our approach follows established practices in network medicine, where higher weights are given to interactions between nodes that are functionally or mechanistically critical ^51,52^.

### 2.2 Drug–MS–EBV Network Construction and Comparative Impact Ranking of MS Immunotherapies

To identify which MS immunotherapies, act broadly across the MS–EBV pathway network (and thus exert higher impact) versus those with narrower effects, we systematically mapped drug–pathway interactions within the network. First, we retrieved the names of FDA- and EMA-approved, currently used (i.e., not discontinued) immunotherapies for MS from DrugBank ^53^ and KEGG ^50^ database. After removing duplicate entries, a total of 33 unique MS immunotherapies were identified. Drug–pathway interactions for these drugs were extracted from KEGG ^41^. In parallel, drug–gene interactions were obtained from DrugBank ^46^, and genes were mapped to KEGG pathways to identify additional drug–pathway interactions relevant to the MS–EBV network. The two datasets were then merged, duplicates were removed, and only drug– pathway interactions involving pathways present in the MS–EBV network were retained. This yielded 81 unique drug–pathway interactions. Of the 33 initial drugs, 17 were found to target 36 pathways within the MS–EBV network.

To quantify the influence of each pathway in the MS–EBV network, we applied the *Disease Node Impact Score* approach as previously described Georgiou et al ^44^. For each pathway (node), the score was defined as the sum of three centrality measures: weighted degree centrality (capturing the strength of direct connections), betweenness centrality (measuring the extent to which a node acts as a bridge in shortest paths), and closeness centrality (reflecting proximity to all other nodes in the network). Calculated using the following equation:

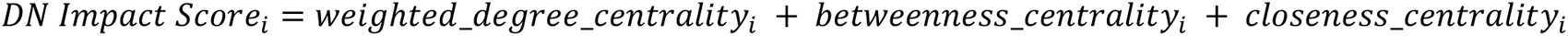

Nodes disconnected from the network were assigned the minimum observed score, consistent with the original method, to avoid excluding potentially relevant but weakly connected pathways. Full details of the algorithm and its rationale are provided in Georgiou et al ^44^.

We then calculated the “Drug Impact Effect Score” for each MS immunotherapy on the MS–EBV network by summing the “Disease Node Impact Scores” of all pathways (nodes) targeted by that drug. This provided a quantitative measure of each drug’s overall influence across the network. Finally, MS immunotherapies were ranked from highest to lowest based on their Drug Impact Effect Score, with higher scores indicating a greater breadth and intensity of impact on the MS–EBV network.

### 2.3 Statistical Comparison of MS Immunotherapies Effects Based on Drug Class in the MS-EBV pathway network

We then compared the relative effects of different MS immunotherapy classes on EBV-associated pathway perturbations in the MS–EBV pathway-pathway network. Therapies were grouped into classes according to their primary mechanism of action (Fumarates, S1P modulators, Interferons, Anti-CD20 mAbs, Anti-CD52, Anti-adhesion [α4-integrin], Glatiramoids, and Glucocorticoid). For each class, we summarized the distribution of individual drug Impact_Scores and ordered classes by their median values. We tested whether Impact_Score differed across classes using a Kruskal–Wallis rank-sum test (non-parametric one-way ANOVA). We reported epsilon-squared (ε^2^) as the effect size with confidence intervals. Given small and imbalanced group sizes (e.g., a single fumarate), we did not perform confirmatory pairwise testing; inference is based on the omnibus Kruskal–Wallis result.

### 2.4 Direct Impact of MS Immunotherapies on Infectious-related pathways in KEGG Database

MS immunotherapies are known to increase the risk of infections or viral reactivation, including EBV reactivation ^29,54–56^. To evaluate this risk mechanistically, we examined the direct effects of these therapies on infectious disease–related pathways. Specifically, we extracted from the drug–MS–EBV pathway network all drug–pathway interactions that mapped to KEGG pathways classified under infectious diseases. Using KEGGREST ^57^, we retrieved the KEGG class assignments for each pathway and further categorized them by pathogen type (viral, bacterial, or parasitic) following KEGG’s annotation. We then constructed a drug–infectious disease pathway interaction subnetwork.

### 2.5 Reconstruction of the Drug–MS–EBV–MS Genetic Variant Association Network and Clustering Analysis

To integrate genetic risk with therapeutic targeting and capture the intersection of MS immunotherapies, EBV-related biology, and host genetics, we reconstructed a drug–MS–EBV–MS genetic variant association network. In this analysis, we excluded drug interactions with infectious disease pathways because these were specifically examined in Section 2.4 to assess infection risk. Here, our focus was restricted to pathways that directly link MS immunotherapies with MS genetic variants embedded in the MS–EBV pathway network, allowing us to disentangle therapeutic effects on host–virus–genetic interactions from those on broader pathogen-related pathways.

MS-associated genetic variants (Concept ID: C0026769) were collected 362 genes from the DisGeNET ^58^ database, and only those variants located in pathways targeted by MS immunotherapies were retained. Since *HLA-DRB1*15:01* allele, the strongest known genetic risk factor for MS^59^, was not captured in the DisGeNET dataset, *HLA-DRB1* was manually added to ensure inclusion of this well-established MS susceptibility locus. Apart from *HLA-DRB1*, no additional variants were manually inserted. The omission of *HLA-DRB1* from DisGeNET likely reflects limitations of the database. From these data, drug–pathway– genetic variant associations were extracted, and a binary presence/absence matrix was created, where each row corresponded to a drug and each column corresponded to a unique MS-associated gene. A value of “1” indicated that the gene was present in at least one pathway targeted by the drug, while “0” indicated no such association.

Jaccard similarity was calculated for all pairwise comparisons between drugs as the number of shared genes divided by the total number of unique genes in either drug. This measure was converted to a Jaccard distance and used as input for hierarchical agglomerative clustering in R. Multiple linkage methods (single, complete, average, ward.D2, mcquitty, median, centroid) were evaluated by calculating the cophenetic correlation coefficient between the input distance matrix and the dendrogram structure, with the method yielding the highest correlation selected for the final analysis.

The optimal number of clusters (*k*) was determined by calculating the average silhouette width for *k* values ranging from 2 to 10, and selecting the value that maximized the mean silhouette score. The final dendrogram was visualized using the factoextra ^60^ package.

### 2.6 Integration of EBV–Host PPIs with MS Variant Genes and Immunotherapy Pathway Targets

We examined whether EBV directly targets proteins encoded by genes associated with MS genetic variants that are also located within pathways modulated by approved MS immunotherapies. Using the collected EBV–host PPI dataset, we isolated EBV interactions involving MS variant–associated genes present in drug-targeted pathways. In parallel, we curated the corresponding set of MS immunotherapies acting on these pathways.

We constructed a tripartite network to illustrate these overlaps, incorporating both drug–MS variant gene associations and EBV protein–MS variant gene interactions. The resulting network contained 37 nodes connected by 72 edges, highlighting the points of convergence between MS therapies, host genetic variants, and EBV proteins.

### 2.7 Assessing Whether MS Immunotherapies Reverse or Reinforce MS–EBV Convergent Transcriptomic Signature

While the comparative impact ranking (Section 2.2) quantified the magnitude of drug effects across the MS–EBV pathway network, it did not capture the *directionality* of these effects. Specifically, it did not establish whether therapies are predicted to counteract or exacerbate EBV-driven transcriptomic signature in MS. To address this, we performed transcriptomics directionality analysis.

We used the LINCS/L1000 collection of signature-level gene sets, specifically the *l1000_cp.gmt* file (chemical perturbations) and the *l1000_aby.gmt* file (antibodies). Each GMT file contains drug-induced gene expression signatures separated into up- and downregulated gene sets. For each therapy (e.g., dimethyl fumarate, rituximab, and ocrelizumab), we curated a synonym list and retrieved all signatures whose names contained any of these synonyms and ended with the labels “… up” or “… down.” This procedure often returned *multiple up and down signatures for a given drug*. To account for this, we merged all up signatures into a single *drug_up* gene set and all down signatures into a single *drug_down* gene set. Duplicate genes were removed so that each therapy was represented by one consolidated up- and one consolidated downregulated signature.

Transcriptomic datasets associated with MS and EBV were retrieved from Enrichr (https://maayanlab.cloud/Enrichr/). For MS, we extracted signatures from the *Disease_Perturbations_from_GEO_up* and *Disease_Perturbations_from_GEO_down* libraries, which contain differential expression results curated from GEO studies and annotated to disease ontologies. We included both human and mouse studies associated with multiple sclerosis (DOID:2377, DOID:2378, C0026769), spanning GEO accessions GSE26484, GSE16461, GSE38010, GSE23832, GSE21942, GSE10064, and GSE842. These datasets capture a broad spectrum of MS-related transcriptomic changes across disease subtypes and experimental platforms.

For EBV, we extracted signatures from the *Microbe_Perturbations_from_GEO_up* and *Microbe_Perturbations_from_GEO_down* libraries. Because human EBV infection datasets are limited, we used transcriptomic perturbations derived from murine gammaherpesvirus 68 (MHV-68), a widely accepted surrogate model for EBV infection ^61,62^. Specifically, we obtained data from GDS4775, which profiles MHV-68 infection across spleen, brain, and liver (microbe IDs 108–113). Together, these datasets provide a proxy for EBV-related immune perturbations.

For both MS and EBV, we merged the upregulated and downregulated gene sets using the same procedure applied to drug perturbation signatures, removing duplicate genes. This resulted in consolidated *MS-up*, *MS-down*, *EBV-up*, and *EBV-down* gene sets. The convergent MS–EBV transcriptomic signature was then defined as the intersection of direction-matched sets: genes upregulated in both MS and EBV (*SigUp*), and genes downregulated in both (*SigDown*). These shared dysregulated modules formed the reference framework for drug reversal and reinforcement analyses.

Each therapy’s *drug_up* and *drug_down* gene sets were compared against the convergent MS–EBV signature. Direction-matched overlaps were quantified in four categories: (i) *SigUp ∩ DrugUp* (concordant up), (ii) *SigDown ∩ DrugDown* (concordant down), (iii) *SigUp ∩ DrugDown* (discordant up), and (iv) *SigDown ∩ DrugUp* (discordant down). These categories yielded a 2×2 contingency table for each drug, representing concordant versus discordant overlaps with the MS–EBV signature.

To statistically evaluate whether a therapy acted primarily as a *reinforcer* (aligning with EBV-associated dysregulation) or a *reverser* (counteracting EBV-driven changes), we applied one-sided Fisher’s exact tests. Enrichment of concordant overlaps (diagonal dominance) indicated reinforcement, whereas enrichment of discordant overlaps indicated reversal. Odds ratios were used to estimate effect sizes, and p-values were adjusted for multiple testing using the Benjamini–Hochberg method. Each drug was then assigned a primary classification (reinforcer, reverser, or neutral) based on the stronger association.

Finally, to aid interpretation, therapies were ranked by −log10(p-value), and the top reinforcers and reversers were visualized in bar plots. This directionality-informed framework complemented the impact ranking by not only quantifying the strength of drug engagement with the MS–EBV network, but also determining whether therapies are predicted to amplify or mitigate EBV-driven transcriptomics signature in MS.

### 2.8 Pathway-Level Enrichment Analysis of the Effects of MS Immunotherapies on the MS–EBV Transcriptomic Signature

To assess the biological pathways modulated by each therapy in respect to the MS–EBV convergent transcriptomic signature, we performed pathway enrichment analysis using KEGG database. Gene symbols were mapped to Entrez IDs with the *org.Hs.eg.db* annotation package (Bioconductor), and enrichment was conducted with the enrichKEGG function from the *clusterProfiler* R package ^63^. The background universe was defined as the union of all genes targeted by any therapy in our dataset.

For each drug, two sets of reversal genes were defined: (i) MS–EBV upregulated genes downregulated by the drug, and (ii) MS–EBV downregulated genes upregulated by the drug. These two gene sets were separately tested for KEGG pathway enrichment against the background universe. Pathways with fewer than five mapped genes were excluded. P-values were adjusted using the Benjamini–Hochberg procedure, and pathways with adjusted *p adjust* < 0.05 were considered significant.

To improve interpretability, results were further filtered to exclude generic or non-informative categories (e.g., “infectious disease,” “substance dependence”, “Immune disease”). Broad KEGG disease pathway classes were removed to avoid redundancy, as they aggregate diverse processes under disease labels without providing mechanistic insight. The analysis focused on pathways representing underlying biological processes rather than broad disease categories, to capture the specific mechanistic effects of MS immunotherapies on the MS–EBV transcriptomic signature. For each drug and reversal set, the statistically significant enriched pathways were identified, then alluvial diagrams, showing the relationships between drugs, KEGG subcategories, and specific enriched pathways were constructed.

## 3. Results

### 3.1 Comparative Impact Ranking of MS Immunotherapies on the MS–EBV Pathway Network

We systematically mapped 33 FDA- and EMA-approved MS immunotherapies onto the MS–EBV pathway network and quantified their relative influence using the Drug Impact Effect Score, representing the summed pathway impact across all targeted nodes and ranked from highest to lowest (see **Figure 2A**). Among these, 17 therapies showed direct engagement with the MS–EBV network, collectively modulating 36 pathways.

**Figure 2.**
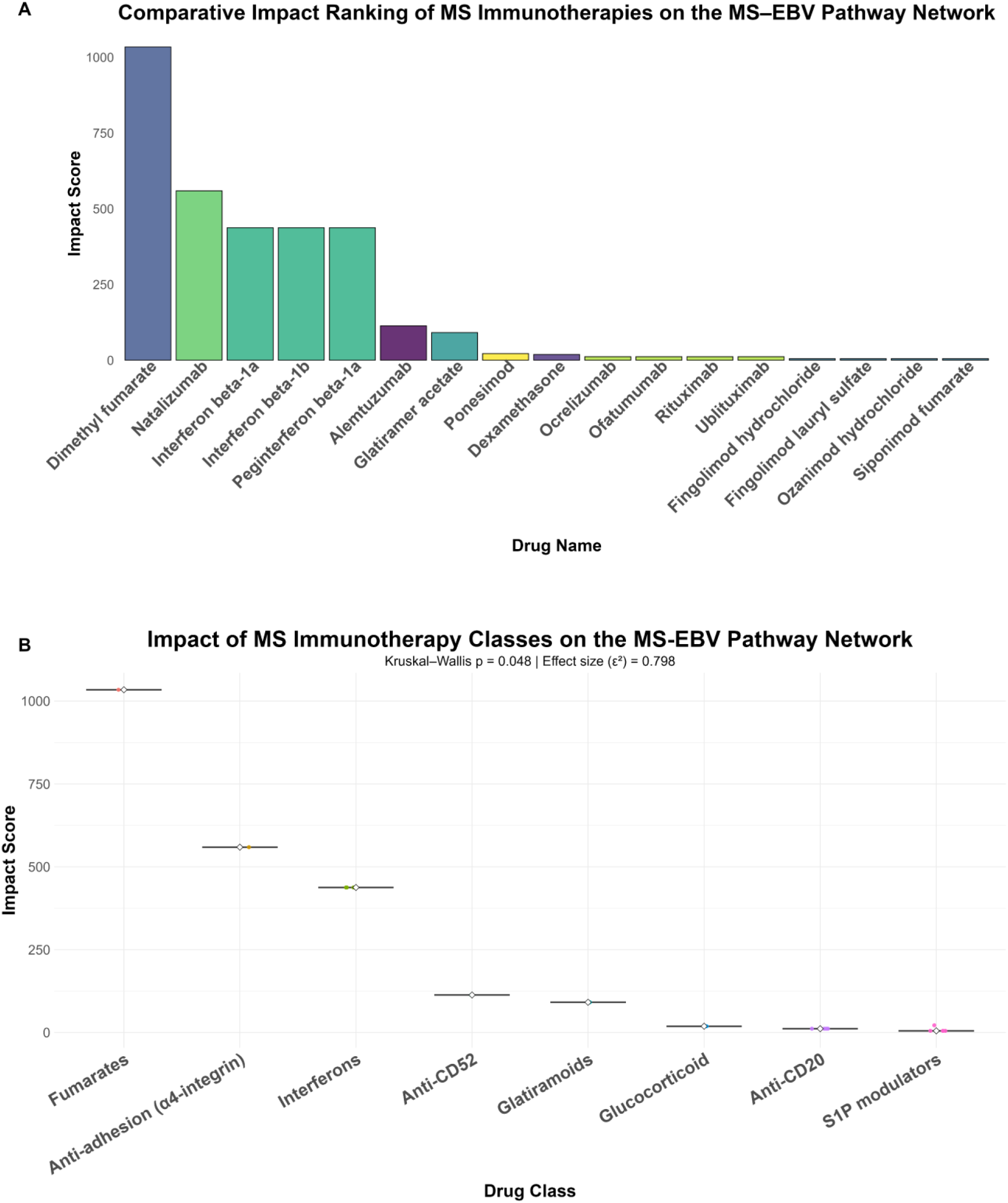
**(A) Comparative Impact Ranking of MS Immunotherapies on the MS–EBV Pathway Network.** Bar plot of Drug Impact Effect Scores for MS immunotherapies, derived from the summed Disease Node Impact Scores of all pathways targeted by each drug. Higher scores correspond to broader and more influential modulation of the MS–EBV pathway network. **(B) Impact of MS Immunotherapy Classes on the MS - EBV Pathway network.** Boxplots show impact scores from the MS–EBV pathway network analysis, grouped by drug class. Horizontal lines indicate medians; jittered points show individual drug scores.

Dimethyl fumarate emerged as the most broadly acting therapy, with the highest overall Drug Impact Effect Score (1034), driven by strong modulation of apoptosis (220.5), NF-κB signaling (136.9), PI3K–Akt signaling (104.3), and Toll-like receptor signaling (106.3) (**Figure 2A**). Natalizumab ranked second (559.2), with effects concentrated on NF-κB signaling (136.9), PI3K–Akt signaling (104.3), and TNF signaling (50.7). Notably, natalizumab also uniquely targeted the intestinal immune network for IgA production (13.1), linking its known efficacy in Crohn’s disease. The interferons (IFN-β1a, IFN-β1b, peg-IFN-β1a) showed nearly identical profiles, each with intermediate Drug Impact Effect Scores (~437). They strongly modulated Toll-like receptor signaling (106.3), PI3K–Akt signaling (104.3), and JAK–STAT signaling (98.3), consistent with their immunomodulatory breadth across innate and adaptive pathways. Alemtuzumab and glatiramer acetate displayed narrower activity (113.3 and 91.3, respectively), mainly targeting phagosome, NK cell-mediated cytotoxicity, and antigen processing/presentation. In contrast, anti-CD20 monoclonal antibodies (ocrelizumab, ofatumumab, rituximab, ublituximab) and S1P modulators (fingolimod, siponimod, ozanimod, ponesimod) had limited influence on the MS–EBV network, with low Drug Impact Effect Scores (<30), reflecting more restricted targeting of hematopoietic lineage or sphingolipid signaling pathways. Taken together, these results reveal that while some therapies (e.g., dimethyl fumarate, natalizumab, interferons) act broadly across the MS–EBV interface, others (e.g., S1P modulators, anti-CD20 antibodies) have narrower effects, potentially contributing to differences in therapeutic efficacy and viral modulation.

### 3.2 MS Immunotherapy Classes Differ in Their Effects on EBV-Associated Pathways in MS

Building on the individual drug ranking, we next assessed whether MS immunotherapies grouped by mechanism of action differ systematically in their modeled effects on MS-EBV network. The network-based analysis revealed clear heterogeneity in the effects of MS immunotherapy classes on EBV-associated pathways (see **Figure 2B**). Fumarates showed the greatest modeled impact, driven entirely by dimethyl fumarate (Impact Score = 1034), which far exceeded all other therapies (**Figure 2B**). The anti-adhesion (α4-integrin) class, represented by natalizumab (559), ranked second in impact. Interferons (interferon beta-1a, interferon beta-1b, peginterferon beta-1a; each 437) followed closely, forming a high-impact group. Intermediate scores were observed for the anti-CD52 class (alemtuzumab, 113), the glatiramoid class (glatiramer acetate, 91.3), and the glucocorticoid class (dexamethasone, 19.0).

By contrast, the anti-CD20 monoclonal antibodies (ocrelizumab, ofatumumab, rituximab, ublituximab; all 11.5) exhibited uniformly low scores. The S1P modulators (fingolimod hydrochloride, fingolimod lauryl sulfate, ozanimod, siponimod, ponesimod; 5.1–21.9) consistently produced the lowest overall class impact, suggesting minimal modulation of EBV-related pathways.

When compared statistically, drug class membership explained a substantial proportion of the variation in Impact Scores (Kruskal–Wallis, df = 7, *p* = 0.048; ε^2^ = 0.798, 95% CI: 0.50–1.00) (**Figure 2B**), confirming significant between-class differences. These findings suggest that certain immunotherapy classes, particularly fumarates, interferons, and natalizumab, may exert stronger indirect effects on EBV-related processes, whereas S1P modulators and anti-CD20 antibodies have relatively limited modeled influence despite their established clinical efficacy in MS.

### 3.3 MS Immunotherapies Directly Target Infectious Disease–Related KEGG Pathways

From the 17 MS immunotherapies identified as targeting pathways within the MS–EBV network, 7 were found to directly target genes associated with infectious disease pathways. To assess the potential infection-related risks of these therapies, we constructed a drug–infectious disease pathway interaction network. The network comprised 35 nodes and 76 edges, including 7 MS drugs, 9 bacterial pathways, 6 parasitic pathways, and 13 viral pathways, including EBV pathway (**Figure 3A**). Of these, four therapies (dimethyl fumarate, interferon beta-1a, interferon beta-1b, and peginterferon beta-1a) were found to directly interact with the EBV pathway.

**Figure 3.**
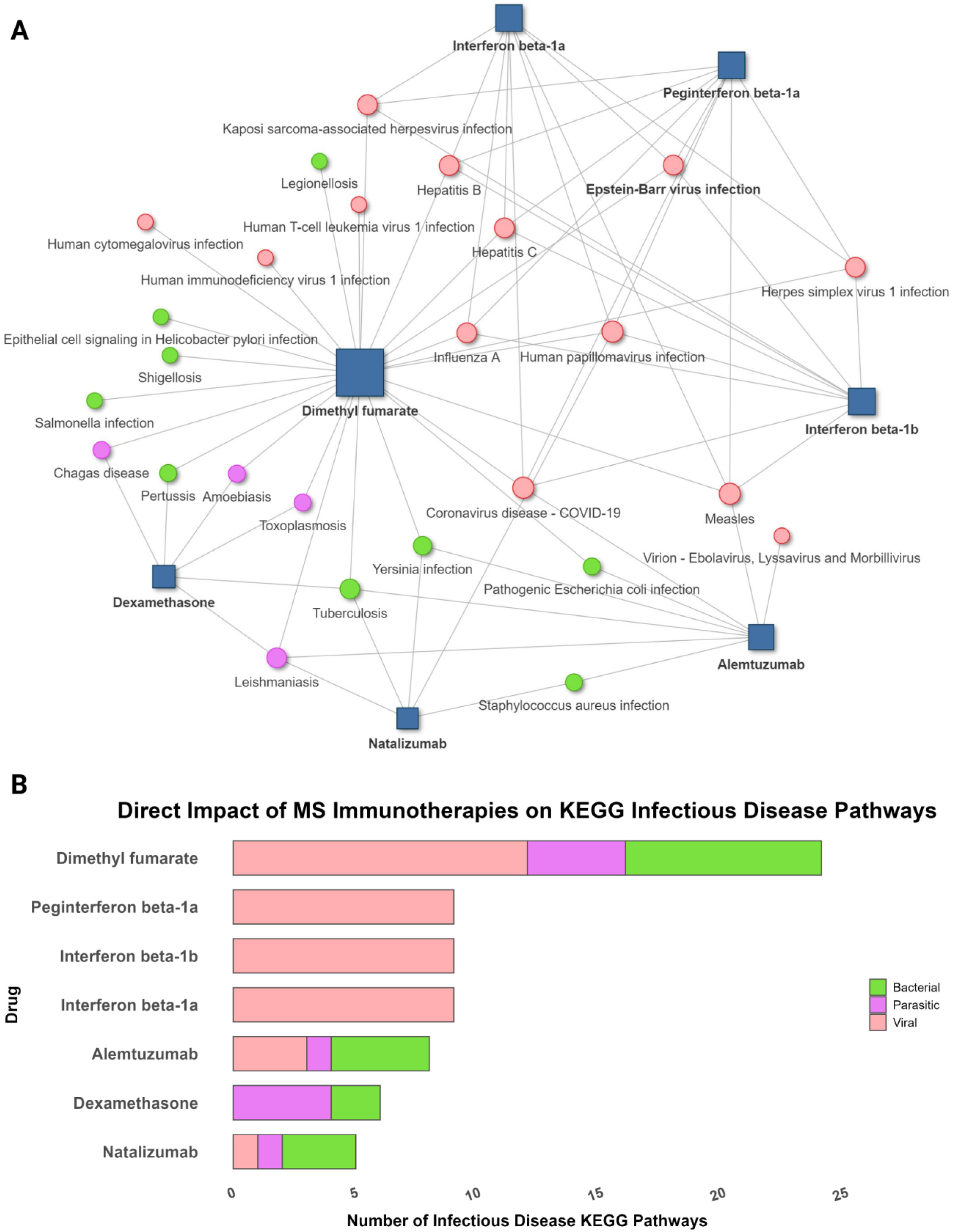
Direct impact of MS immunotherapies on infectious disease–related KEGG pathways. **(A)** Drug–infectious disease pathway interaction network. Blue squares represent MS immunotherapies, while circles represent KEGG infectious disease pathways (red = viral, green = bacterial, purple = parasitic). The EBV pathway is highlighted in bold. **(B)** Barplot summarizing the number of infectious diseases KEGG pathways directly targeted by each of the seven MS immunotherapies.

Analysis of the seven MS immunotherapies that directly targeted infectious disease pathways revealed distinct patterns of connectivity. Dimethyl fumarate showed the broadest impact, targeting 24 infectious disease pathways in total, including 8 bacterial, 4 parasitic, and 12 viral pathways (**Figure 3B**). Notably, dimethyl fumarate was linked to multiple viral pathways (e.g., influenza A, HSV-1, human cytomegalovirus, HIV, hepatitis B/C, and EBV), bacterial pathways (e.g., *Salmonella*, *E. coli*, *Shigella*), and parasitic pathways (e.g., leishmaniasis, amoebiasis), underscoring its pleiotropic effects (**Figure 3A**).

Natalizumab also displayed broad connectivity, targeting 5 pathways (3 bacterial, 1 parasitic, and 1 viral), including viral infections (HPV), parasitic diseases (leishmaniasis), and bacterial infections (*Yersinia*, *Staphylococcus aureus*). Alemtuzumab targeted 8 pathways, spanning 4 bacterial, 1 parasitic, and 3 viral, while dexamethasone was associated with 6 pathways (2 bacterial and 4 parasitic).

In contrast, the interferons exhibited highly selective effects, mapping exclusively to viral pathways. Interferon beta-1a, interferon beta-1b, and peginterferon beta-1a each interacted with 9 viral pathways, including influenza A, COVID-19, herpes simplex virus 1, hepatitis B/C, and EBV.

Dexamethasone targeted six infectious disease pathways: four parasitic (Toxoplasmosis, Amoebiasis, Chagas disease, and Leishmaniasis) and two bacterial (Pertussis and Tuberculosis). In contrast, Alemtuzumab engaged eight infectious disease–related pathways, including four bacterial (E. coli infection, Staphylococcus aureus infection, Yersinia infection, and Tuberculosis), three viral (e.g COVID-19, Measles), and one parasitic (Leishmaniasis).

Together, these findings suggest that the direct targeting of infectious disease pathways by MS immunotherapies may help explain their variable associations with infection and viral reactivation risks observed in clinical practice. In particular, the observation that several therapies directly interact with EBV-related pathways highlights a potential mechanism by which these drugs may influence EBV-driven processes in MS pathogenesis.

### 3.4 MS Immunotherapy Interactions with MS Genetic Variants in the MS–EBV Pathway Network

To investigate the possible interaction of MS immunotherapies with genetic variants linked to MS, for each therapy we identified MS-associated genetic variant genes contained inside the pathways they target. For this part of the analysis, we excluded infectious disease pathways targeted by the drugs, as the aim was to focus on biological pathways directly relevant to MS pathogenesis and EBV-related processes. This approach allowed us to assess how immunotherapies overlap with host genetic risk within immune signaling, antigen presentation, and other MS-relevant pathways, rather than confounding effects from pathogen-related mechanisms.

This analysis identified substantial heterogeneity in variant coverage across therapies (**Figure 4A,B**). Dimethyl fumarate and Natalizumab exhibited the broadest footprint, interacting with 52 MS variant-associated genes across diverse immune-regulatory pathways. Among all therapies, the S1P receptor modulators fingolimod (hydrochloride and lauryl sulfate), ozanimod, and siponimod involved the fewest MS-associated variants, with only three variants identified for each drug.

**Figure 4.**
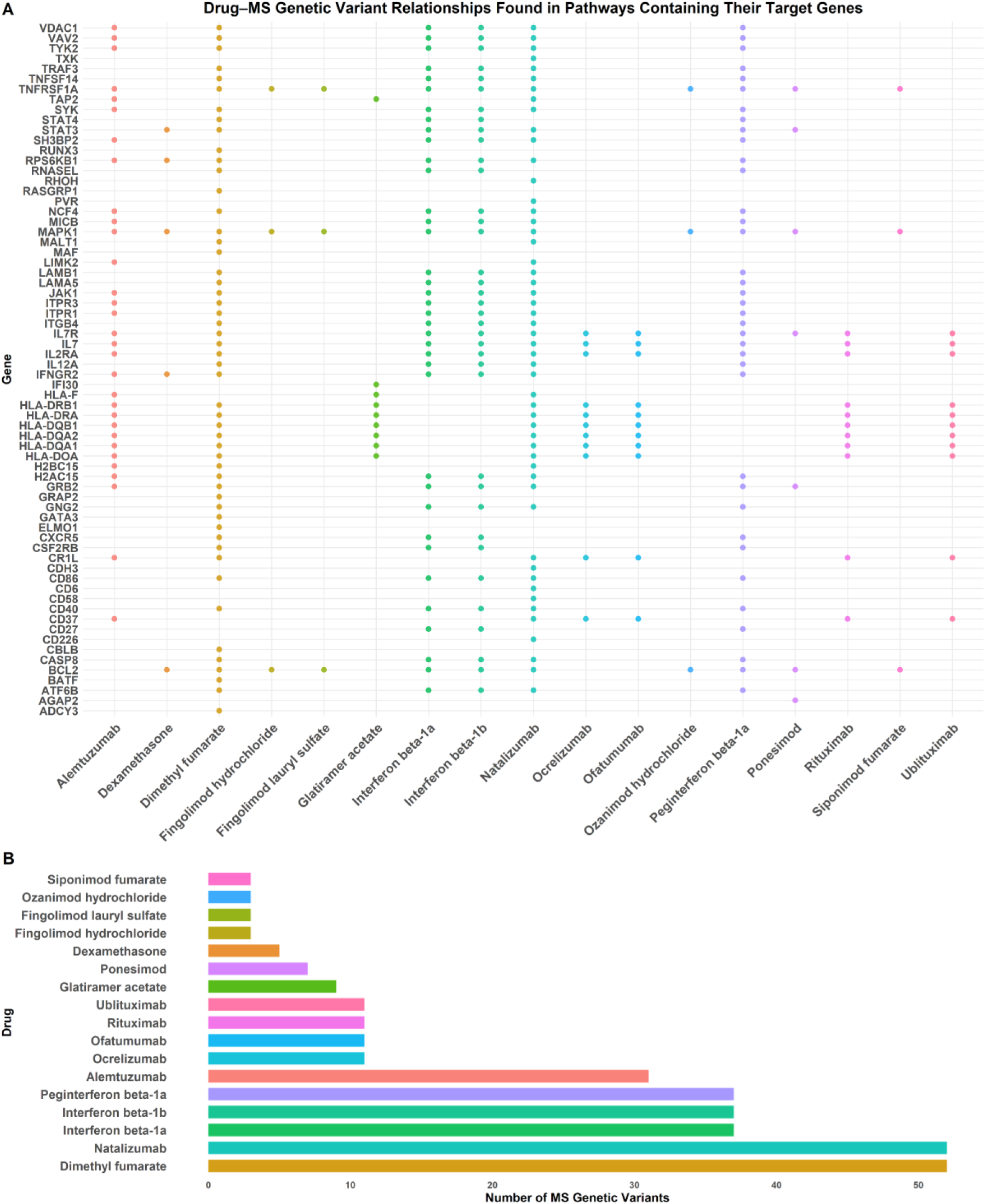
Interaction of MS immunotherapies with MS genetic variants through biological pathways. **(A)** Dot plot showing the overlap between MS genes-associated genetic variants and immunotherapies. **(B)** Bar graph summarizing the total number of MS genetic variants associated with each drug.

Dimethyl fumarate intersected with pathways containing *HLA-DRB1* (the strongest genetic risk factor for MS), along with *IL7R* and *TYK2*, both key immune regulators previously implicated in MS susceptibility. These genes were embedded within major immune pathways, including NF-κB, PI3K–Akt, Th1/Th2 differentiation, and T-cell receptor signaling, underscoring the convergence of this therapy with fundamental host–genetic and immune mechanisms of MS.

Natalizumab was associated with a wide range of MS genetic variants through the pathways it engages. These included strong antigen presentation genes such as *HLA-DRB1, HLA-DQA1, HLA-DQB1,* and *HLA-DRA* within phagosome-related pathways, reinforcing overlap with the strongest known MS risk factor. Variants in *IL7R* and *TYK2* were also present in pathways linked to natalizumab’s targets (PI3K–Akt and osteoclast differentiation), alongside additional immune regulators such as *CD40, TNFRSF1A, STAT3,* and *CASP8*.

Interferon therapies (interferon beta-1a, beta-1b, and peginterferon beta-1a) collectively overlapped with 37 MS-associated genetic variants through the pathways they engage. Prominent among these were *IL7R, IL2RA, and IL7* within cytokine–cytokine receptor interaction and PI3K–Akt signaling pathways, all central immune regulators with established roles in MS susceptibility. Variants in *TYK2* and *STAT4* were also represented within necroptosis pathways, highlighting convergence on JAK–STAT signaling. Additional associations included *TNFRSF1A* and *CD40* within cytokine pathways, *STAT3* and *CASP8* in necroptosis, and antigen presentation–related genes such as *CD86* and *TRAF3* in toll-like receptor signaling.

The strongest MS risk allele, *HLA-DRB1*, was found within pathways connected to multiple classes of immunotherapies. Alemtuzumab and natalizumab converged on *HLA-DRB1* through phagosome-mediated antigen presentation, while glatiramer acetate involved *HLA-DRB1* in antigen processing and presentation. Dimethyl fumarate linked to HLA-DRB1 within Th1/Th2 differentiation pathways, highlighting its role in shaping adaptive immune responses. Importantly, all four anti-CD20 therapies (ocrelizumab, ofatumumab, rituximab, and ublituximab) also overlapped with *HLA-DRB1* via the hematopoietic cell lineage pathway, reflecting its central role in B cell biology. Together, this demonstrates that *HLA-DRB1* is a common genetic intersection point across distinct therapeutic classes including T cell targeting therapies such as alemtuzumab, glatiramer acetate, dimethyl fumarate and natalizumab, and B cell depleting therapies such as the anti-CD20 monoclonal antibodies, underscoring its fundamental contribution to MS pathogenesis.

Dimethyl fumarate, alemtuzumab, glatiramer acetate, and natalizumab target pathways that are shared between MS and EBV in the MS-EBV pathway network, linking MS susceptibility genes to EBV-associated immune processes. Alemtuzumab and natalizumab converge on the phagosome pathway enriched for HLA class II variant genes (*HLA-DRB1, HLA-DQA1, HLA-DQB1, HLA-DRA, TAP2*), central to both MS risk and EBV antigen display. Dimethyl fumarate and natalizumab further intersect with NF-κB signaling, which contains MS variants such as *CD40*, *TRAF3, TNFRSF1A*, and *MALT1*, key regulators of EBV latency and immune evasion. Natalizumab also engages the lipid and atherosclerosis pathway, which includes the *IL12A* MS variant and represents a shared pathogenic axis between MS and EBV. This is biologically relevant, as EBV manipulates host lipid metabolism during latent and lytic infection, while dysregulated lipid pathways, particularly cholesterol and phospholipids, are implicated in MS pathogenesis. Together, these results show that several MS therapies converge not only on general immune signaling but also on pathways simultaneously implicated in MS risk and EBV biology, reinforcing a genetic and functional bridge between therapeutic mechanisms and host–virus interactions.

In summary, this analysis highlights that MS immunotherapies engage genetic risk factors embedded within pathways central to both MS pathogenesis and EBV biology. The repeated convergence on HLA-DRB1 and other immune regulators underscores their role as unifying genetic nodes across therapies, while the targeting of NF-κB and lipid metabolism pathways further reveals shared mechanisms linking host genetics, viral biology, and therapeutic action. These findings suggest that MS drug effects may be understood not only in terms of immune modulation, but also in their alignment with genetic and viral determinants of disease, providing a genetic–viral framework for interpreting therapeutic impacts.

### 3.5 Clustering of MS Immunotherapies by Overlap in Genetic Variant Associations

To compare MS immunotherapies based on shared genetic variant associations, we performed hierarchical agglomerative clustering using Jaccard distance between drug–gene relationships. Among the linkage methods tested, the average method produced the highest cophenetic correlation coefficient (0.978), and was therefore selected for the final analysis. The optimal partitioning yielded six distinct clusters (**Figure 5**).

**Figure 5.**
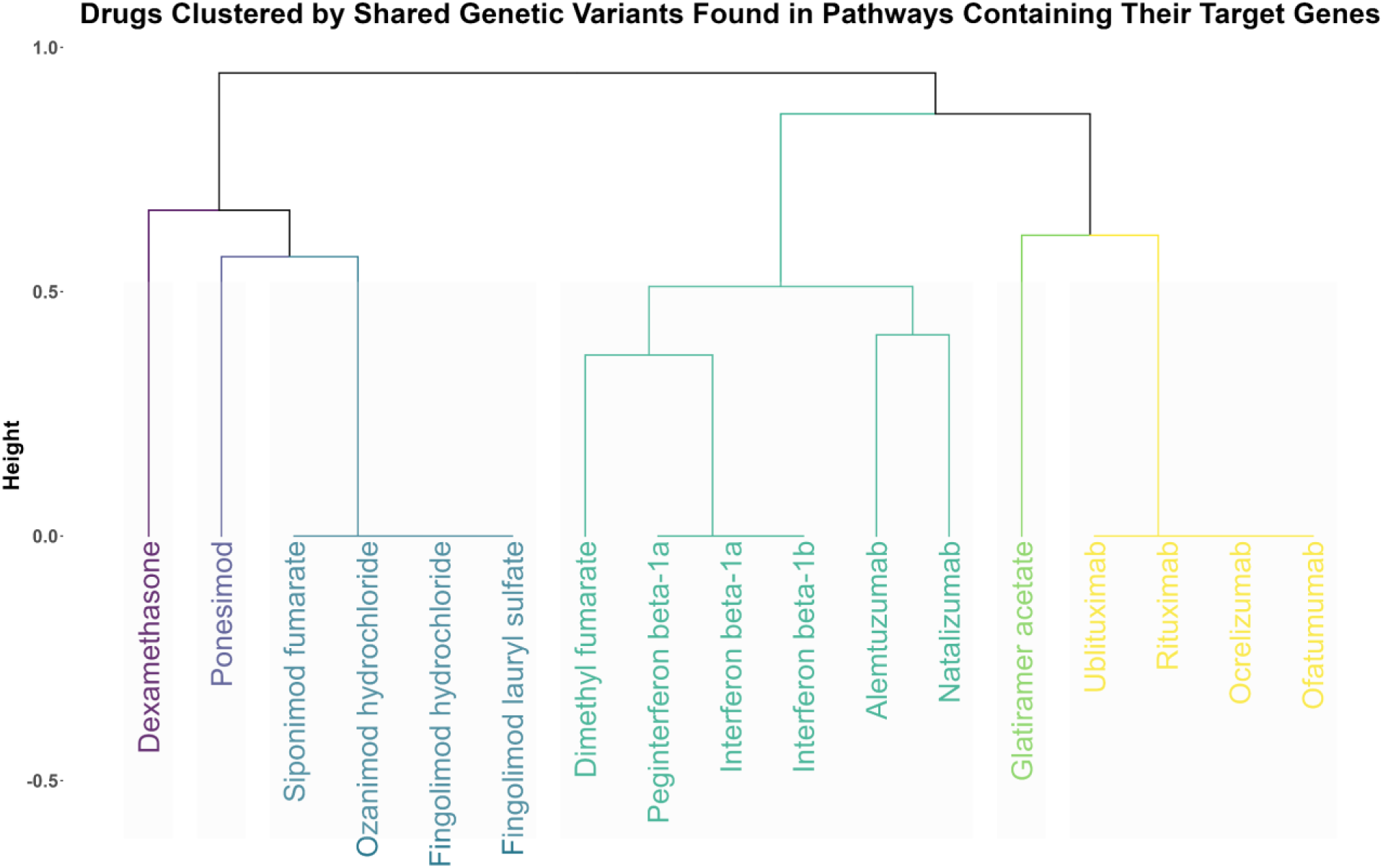
Hierarchical clustering of drug–gene associations. Clustering of MS immunotherapies based on overlap in MS-associated genetic variants contained within biological pathways reveals distinct therapeutic groupings.

The clustering recapitulated both known pharmacological classes and unexpected genetic overlaps. Dexamethasone formed its own cluster, positioned at one extreme of the dendrogram, while the anti-CD20 therapies (ocrelizumab, rituximab, ofatumumab, and ublituximab) grouped tightly at the opposite end, reflecting their distinct variant profiles. S1P receptor modulators (fingolimod, ozanimod, and siponimod) clustered together, consistent with their shared mechanism. Interestingly, interferons clustered with dimethyl fumarate, alemtuzumab, and natalizumab, revealing convergence on overlapping MS-risk variants despite their mechanistic differences. Finally, glatiramer acetate formed an independent cluster, highlighting its unique genetic association footprint. The results revealed both alignment with known drug classes and unexpected overlaps, indicating shared genetic footprints among otherwise distinct mechanisms. Together, this shows that MS therapies can be grouped not only by mechanism of action but also by their overlap with host genetic risk, suggesting a genetic basis for both class-specific and cross-class therapeutic effects.

### 3.6 EBV Proteins Target MS Variant Genes contained in Pathways Targeted by Immunotherapies

To determine whether EBV directly targets MS variant–associated genes that lie within pathways modulated by MS immunotherapies, we constructed a tripartite network integrating drug–pathway associations, MS variant genes, and EBV protein–host interactions. The final network comprised 37 nodes and 72 edges, capturing the overlap among MS therapies, variant-associated human genes, and EBV proteins. While MS immunotherapies do not directly bind or inhibit MS variant genes themselves, they target immune pathways in which these variant genes play critical roles. EBV, through its proteins, was found to interact with several of MS variant–associated genes, highlighting points of convergence (**Figure 6**).

**Figure 6.**
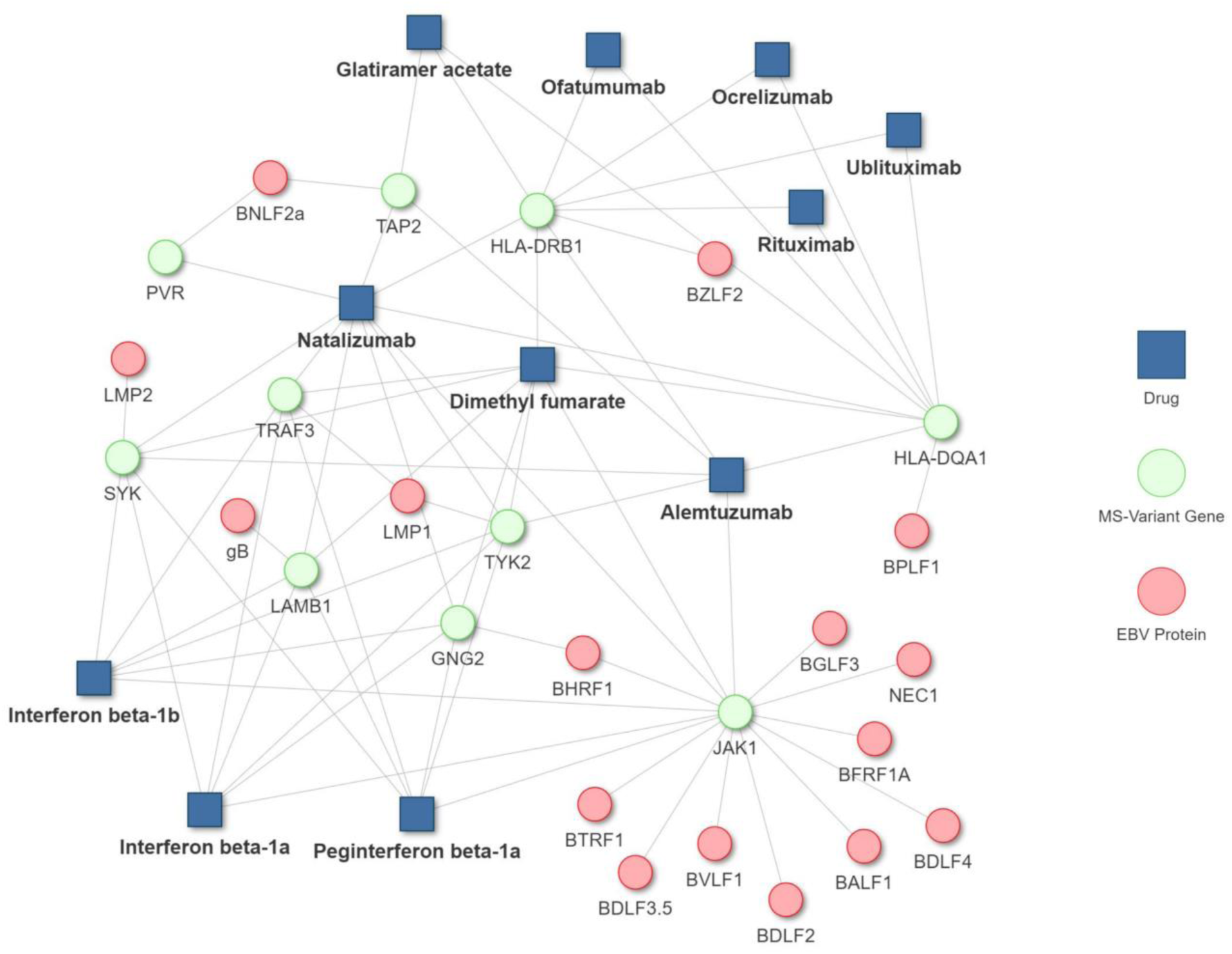
Tripartite network of EBV proteins, MS variant–associated genes, and MS immunotherapies. The network depicts converging relationships between EBV proteins (red), MS variant–associated genes (green), and MS immunotherapies (blue).

The analysis showed that JAK1, a gene associated with MS variants, is targeted by multiple EBV proteins (BDLF4, BALF1, BTRF1, BVLF1, BHRF1, BGLF3, BDLF2, BDLF3.5, BFRF1A, NEC1). MS drugs such as Alemtuzumab, Dimethyl fumarate, Interferons, and Natalizumab modulate signaling pathways including JAK-STAT, PI3K-AKT, Toll-like receptor, and Th1/Th2/Th17 differentiation pathways in which JAK1 plays a central role.

Similarly, SYK, engaged by the EBV protein LMP2, participates in immune signaling cascades that are modulated by Dimethyl fumarate, Interferons, Alemtuzumab, and Natalizumab. TYK2, targeted by EBV LMP1, is another critical mediator of cytokine signaling and immune regulation pathways influenced by these therapies. EBV also directly interacts with TRAF3 via LMP1. TRAF3 is a key regulator of NF-kB, IL-17, and TNF signaling pathways, which are modulated by Dimethyl fumarate, Interferons, Alemtuzumab, and Natalizumab.

HLA-DQA1 and HLA-DRB1 are targeted by the EBV proteins BPLF1 and BZLF2, respectively. These genes function within antigen presentation and hematopoietic cell lineage pathways that are influenced by therapies such as Alemtuzumab, Glatiramer acetate, Dimethyl fumarate, Natalizumab, and anti-CD20 antibodies (Ocrelizumab, Ofatumumab, Ublituximab, Rituximab). In addition, the structural and adhesion molecule LAMB1, a variant-associated gene, is targeted by the EBV gB protein. LAMB1 is positioned in the PI3K-Akt signaling pathway, which is also modulated by Dimethyl fumarate, Interferons, and Natalizumab.

Taken together, these results demonstrate that EBV proteins and MS therapies converge on overlapping immune pathways through distinct routes: EBV proteins directly interact with host variant-associated genes, whereas MS therapies modulate the broader pathways in which these genes operate. This convergence suggests that EBV manipulation of host processes may interfere with, or alternatively be counterbalanced by, therapeutic mechanisms, underscoring a complex interplay between viral reactivation, genetic susceptibility, and treatment response in MS. These findings provide mechanistic clues for designing adjunct therapeutic strategies that pair immune modulation with EBV-targeted interventions to enhance treatment efficacy and disease control.

### 3.7 Classification of MS Immunotherapies as Reinforcers or Reversers of the Convergent MS–EBV Transcriptomic Signature

To directly assess whether MS immunotherapies reinforce or reverse the convergent MS–EBV transcriptomic signature, we applied a comparative framework contrasting drug-induced transcriptomic responses against the MS–EBV overlap signature (Methods 2.7). Each therapy was evaluated for concordant (reinforcing) versus discordant (reversing) effects relative to the joint up- and down-regulated gene sets.

Across the 15 therapies for which high-quality perturbational signatures were available, results revealed a clear stratification into two categories (**Figure 7**). Reinforcers included dimethyl fumarate, dexamethasone, dalfampridine, mitoxantrone, fingolimod, triamcinolone, and interferon-β1b, all of which showed transcriptomic responses aligned with the convergent MS–EBV signature. Reversers comprised cortisol, prednisolone, teriflunomide, hydrocortisone, cladribine, methylprednisolone, interferon-β1a, and rituximab, which induced gene expression shifts opposite to the MS–EBV overlap, suggestive of potential antagonism of shared pathogenic processes.

**Figure 7.**
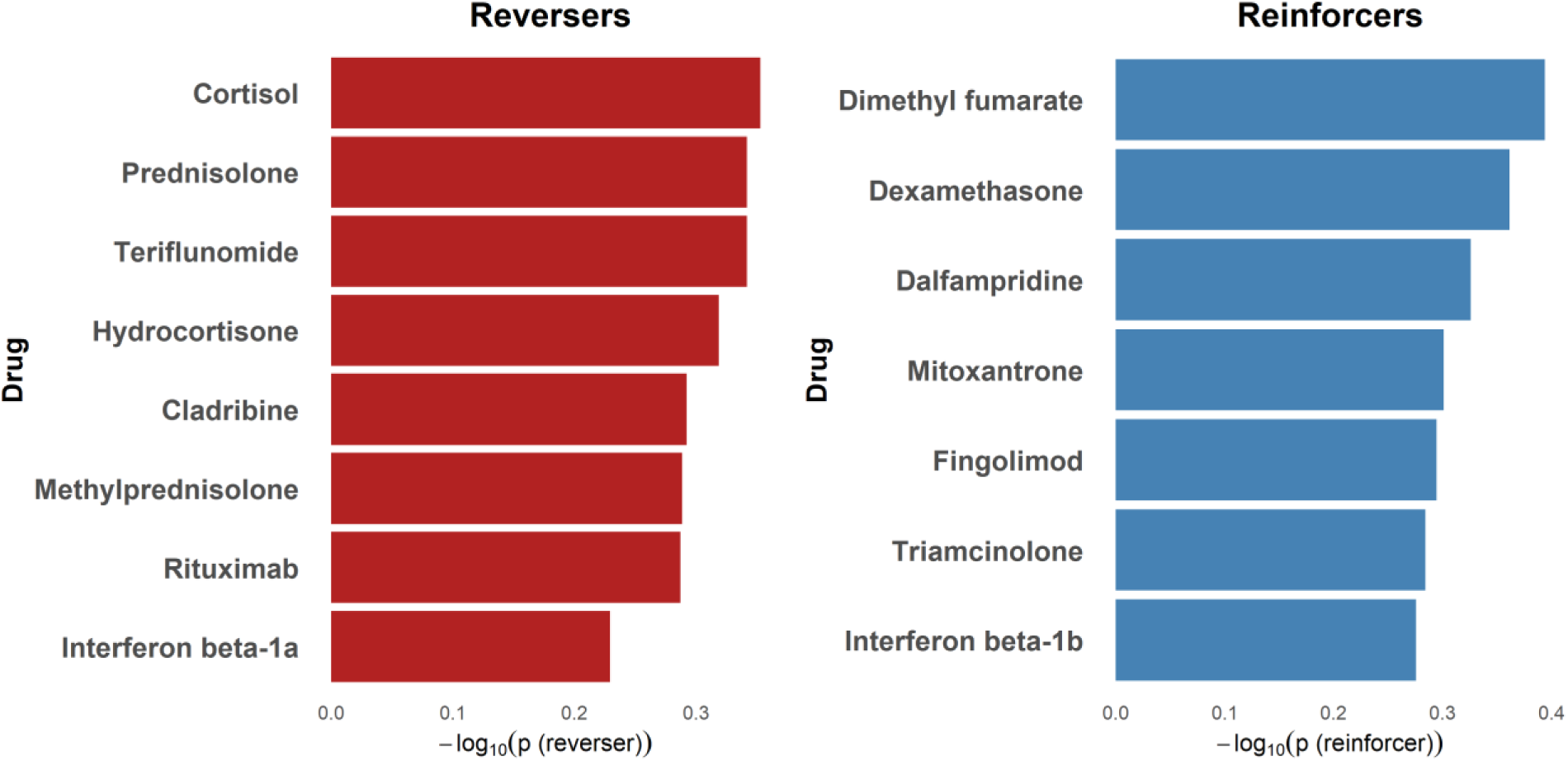
Classification of MS immunotherapies as reinforcers or reversers of the convergent MS–EBV transcriptomic signature.

Notably, corticosteroids split across both groups: dexamethasone and triamcinolone clustered as reinforcers, whereas prednisolone, hydrocortisone, methylprednisolone, and cortisol were classified as reversers. Similarly, the two interferon-β isoforms diverged, with IFN-β1b classified as a reinforcer and IFN-β1a as a reverser.

Although none of the associations reached statistical significance after multiple-testing correction, the reinforcer/reverser assignments recapitulated expected biological behaviors, supporting the validity of this framework as a hypothesis-generation tool.

Overall, the reinforcement–reversal analysis underscores that drugs currently grouped under the same clinical class may exert fundamentally distinct transcriptomic influences on the convergent MS–EBV signature. These findings suggest that transcriptome-informed classification could refine therapeutic strategies, particularly in EBV-seropositive MS patients, by identifying candidates that oppose rather than reinforce EBV-driven immune perturbations.

### 3.8 Pathway-Level Reversal by MS Immunotherapies of the MS–EBV Convergent Transcriptomic Signature

#### 3.8.1 Downregulation of MS–EBV Upregulated Transcriptomic Signature

Pathway analysis of reversal effects against the MS–EBV convergent transcriptomic signature revealed modulation of cytokine signaling pathways (**Figure 8A**). Thirteen therapies (cladribine, dimethyl fumarate, fingolimod, hydrocortisone, interferon-β1a, interferon-β1b, methylprednisolone, mitoxantrone, prednisolone, rituximab, teriflunomide, and triamcinolone) consistently downregulated the KEGG pathway *“Cytokine–cytokine receptor interaction,”* which was pathologically upregulated in the MS–EBV overlap transcriptomic signature. This pathway represents the host cytokine and receptor signaling network, a key immune axis dysregulated by EBV infection and MS inflammation.

**Figure 8:**
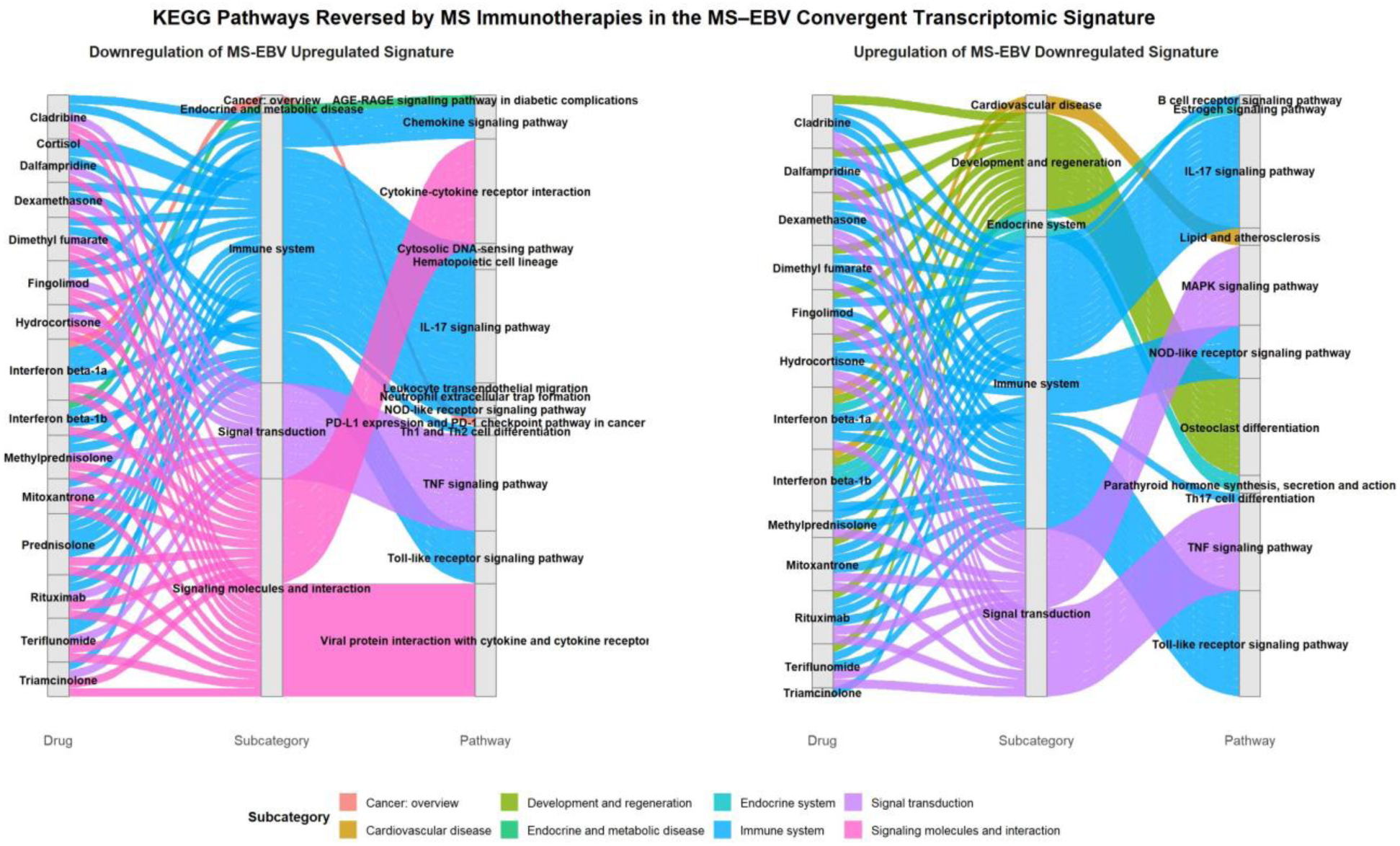
Alluvial plots show drug–subcategories-pathway relationships identified by KEGG enrichment of reversal gene sets. **(A)** Pathways downregulated by therapies within the MS–EBV upregulated signature. **(B)** Pathways upregulated by therapies within the MS–EBV downregulated signature

Interestingly, the enrichment analysis also revealed that 13 drugs (cladribine, dalfampridine, dexamethasone, dimethyl fumarate, fingolimod, hydrocortisone, interferon-β1a, methylprednisolone, mitoxantrone, prednisolone, rituximab, teriflunomide, and triamcinolone)) also reversed the *“Viral protein interaction with cytokine and cytokine receptor,”* pathway which involves viral strategies to evade immune surveillance through cytokine mimicry, homologs, and soluble cytokine-binding proteins. Together, these findings suggest that a broad range of MS therapies, despite diverse pharmacological classes, converge on suppressing EBV-exploited cytokine signaling pathways. By dampening the hyperactive cytokine–cytokine receptor network and counteracting viral cytokine mimicry, these agents may mitigate a shared immunopathogenic mechanism linking EBV persistence with MS progression.

Additional pathways reversed included IL-17 signaling TNF signaling, chemokine signaling, Toll-like receptor signaling, and NOD-like receptor signaling. These innate and adaptive immune pathways are central to both MS pathogenesis and EBV persistence. For example, cladribine, dimethyl fumarate, fingolimod, and corticosteroids consistently reversed IL-17 and TNF pathway enrichment, both implicated in chronic inflammation and EBV latency control.

#### 3.8.2 Upregulation of MS–EBV Downregulated Transcriptomic Signature

Complementary to these effects, many therapies upregulated pathways suppressed in the MS–EBV overlap signature (**Figure 8B**). Notably, 9 therapies (cladribine, dalfampridine, dexamethasone, fingolimod, hydrocortisone, Methylprednisolone, Mitoxantrone, Rituximab, Teriflunomide) upregulated the MAPK signaling which might reflect restoration of pathways abnormally suppressed in the MS–EBV convergent signature, indicating normalization of host defense and homeostatic modules rather than pro-inflammatory activation.

Across multiple therapies (cladribine, dalfampridine, dexamethasone, dimethyl fumarate, fingolimod, hydrocortisone, interferon-β1a, interferon-β1b, mitoxantrone, rituximab, and teriflunomide), we observed upregulation of the osteoclast differentiation pathway, a process suppressed in the convergent MS–EBV signature. This suggests that immunotherapies may restore signaling processes governing osteoclastogenesis, which are abnormally dampened in MS–EBV overlap. Similarly, multiple therapies enhanced IL-17 signaling, TNF signaling, and Toll-like receptor signaling, pathways essential for immune activation and viral sensing.

#### 3.8.3 Mixed Directional Effects within the Same Pathway

Interestingly, several therapies demonstrated bidirectional regulation of the same pathways, appearing both in the downregulation of MS–EBV upregulated signatures and in the upregulation of MS–EBV downregulated signatures. This pattern was most evident in several key inflammatory pathways, including IL-17, TNF, and Toll-like receptor. The IL-17 pathway was reversed in both directions, with agents such as cladribine, dimethyl fumarate, fingolimod, corticosteroids, and rituximab contributing to both downregulation of MS–EBV upregulated genes and upregulation of MS–EBV downregulated genes. TNF signaling displayed a similar bidirectional profile, being influenced in both directions by cladribine, dexamethasone, fingolimod, hydrocortisone, mitoxantrone, and rituximab. Toll-like receptor signaling also appeared in both categories, reversed dalfampridine, dexamethasone, dimethyl fumarate, and rituximab.

## 4. Discussion

This study is the first to systematically apply a virus-informed network pharmacology framework to assess how approved MS immunotherapies intersect with EBV-driven disease mechanisms. By integrating MS-EBV pathway interaction, EBV–host-MS interactomes, MS genetic risk loci, MS-EBV transcriptomic signatures and drug-induced transcriptomic responses, we show that therapies differ markedly in their capacity to engage viral pathways, intersect with susceptibility alleles, and reshape EBV-related expression signatures. Our results reveal that approved therapies differentially engage EBV-driven pathways, converge on genetic risk nodes such as HLA-DRB1, and stratify into “reinforcers” or “reversers” of MS-EBV convergent transcriptomic signatures. Some agents reinforce transcriptomic patterns associated with EBV persistence, while others act in the opposite direction, reversing convergent MS–EBV modules. These distinctions provide mechanistic context to the “immunosuppression paradox” in MS: drugs developed to suppress autoimmunity do not uniformly influence EBV biology, and in some cases may inadvertently support the viral processes that sustain disease. Unlike prior experimental or clinical approaches, which are typically limited to single therapies or models, this systems-level analysis provides a broad comparative framework across multiple approved MS drugs, yielding novel biological insights into the virus–host–therapy interface and paving the way for precision, virus-informed treatment strategies in MS.

The advantage of these approaches is that comparative network pharmacology can investigate a pool of drugs across conditions simultaneously. Experimental approaches such as cell-based assays, animal models, or human trials are limited in their ability to comprehensively evaluate how multiple therapies interact with EBV-driven mechanisms in MS. Cell-based systems lack the ability to capture system-level interactions or the full complexity of MS pathogenesis, while mouse models require a separate experimental setup for each therapy, making a comparative analysis across 15–20 drugs prohibitively time-consuming and resource-intensive. Human studies are further constrained by ethical considerations, patient heterogeneity, and the impracticality of systematically testing all therapies. Network pharmacology provides a scalable alternative, enabling simultaneous, systems-level comparison of all approved drugs within the EBV–MS framework. Within this systems framework, we compared all approved MS therapies to reveal how their molecular footprints converge or diverge at the EBV–host-MS interface.

Dimethyl fumarate emerged as the most influential therapy across the MS–EBV network, modulating apoptosis, NF-κB, PI3K–Akt, and Toll-like receptor signaling, pathways central to both MS immunopathogenesis and EBV latency. EBV exploits NF-κB via its latent membrane protein 1 (LMP1) ^64^. Natalizumab ranked second in overall network impact, strongly modulating NF-κB, PI3K–Akt, and TNF signaling pathways. Serological profiling during natalizumab therapy has revealed increases in anti-VCA IgG, consistent with EBV reactivation ^29^. Together, the network and serological evidence suggest that while natalizumab engages multiple EBV-related immune pathways, its potent immunomodulatory action, by restricting immune cell trafficking, may impair antiviral surveillance, allowing subclinical EBV reactivation rather than suppression. Our analysis also identified a unique effect of natalizumab on the intestinal immune network for IgA production, aligning with its established efficacy in Crohn’s disease. However, despite its effectiveness in both MS and Crohn’s disease, natalizumab carries a known risk of John Cunningham virus (JCV)–driven progressive multifocal leukoencephalopathy (PML), a rare but often fatal demyelinating condition ^54,65^. Although IgA is not the primary defense against JCV, disruption of mucosal immune networks and altered B-cell trafficking may provide a mechanistic link to JCV pathogenesis, given that B cells serve as a viral reservoir. Clinically, natalizumab’s immune exclusion mechanism underscores the trade-off between effective CNS immune control and impaired systemic viral surveillance, consistent with its well-documented association with JCV-driven PML ^54,65^.

Interferon-β therapies ranked third in overall impact across the MS–EBV pathway network, strongly engaging pathways such as JAK–STAT and Toll-like receptor signaling pathways. Functionally, IFN-β reduces T-cell responses to EBNA1 ^66^, potentially dampening autoimmune mimicry but also weakening antiviral immunity. EBV further undermines interferon activity through its BZLF1 protein, which suppresses IFN-γ receptor expression and downstream signaling ^67^. Notably, while pathway-level analysis indicated highly similar targeting across IFN-β isoforms, based on collected drug–pathway and drug–gene interactions, our transcriptomic directionality analysis revealed divergent effects with IFN-β1a behaving as a reverser of the MS–EBV convergent transcriptomic signature, whereas IFN-β1b acting as a reinforcer. This discrepancy may reflect several mechanisms. First, the isoforms could engage context-dependent or unannotated targets not represented in pathway databases. Second, despite convergence on canonical pathways such as JAK–STAT and Toll-like receptor signaling, biochemical differences between IFN-β1a and IFN-β1b, including glycosylation status, receptor binding affinity, and pharmacokinetics, may alter the strength, duration, or balance of downstream signaling, producing distinct transcriptomic signatures. Third, cell type–specific responses or differences in secondary network effects may also contribute. Collectively, these factors highlight how subtle isoform-specific differences can translate into divergent transcriptomic outcomes and may underlie variability in therapeutic responses to interferon in EBV-positive MS patients.

S1P modulators (e.g., fingolimod) exhibited weak MS-EBV network engagement. Mechanistically, fingolimod sequesters lymphocytes in lymphoid tissue ^31^, limiting CNS infiltration but also impairing systemic antiviral surveillance. This has clinical consequences since fingolimod-treated individuals show poor seroconversion after SARS-CoV-2 vaccination ^32^, and rare cases of EBV-positive CNS lymphomas and EBV+ B cell lymphomas have been reported ^33^. Our network findings are in line with these observations, showing minimal EBV-directed network reversal, consistent with the notion that S1P modulators may inadvertently create niches for viral persistence.

Anti-CD20 antibodies (rituximab, ocrelizumab, ofatumumab, ublituximab) showed relatively narrow EBV network effects. Yet they directly deplete EBV-harboring memory B cells, the main viral reservoir. Ocrelizumab reduces proliferation to EBV-infected lymphoblastoid cell lines and dampens IFN-γ responses to EBV antigens ^37^, while rituximab is frontline therapy for EBV-driven lymphoproliferative disorders^39^. It has been proposed that their strong efficacy in MS derives primarily from elimination of EBV-infected memory B cells ^38^. Our classification of rituximab as a “reverser” of MS–EBV transcriptomic overlap supports this antiviral reservoir-depletion hypothesis.

Taken together, these drug-specific findings reveal a continuum of antiviral versus immunosuppressive network behaviors. Dimethyl fumarate and anti-CD20 antibodies broadly opposed EBV-driven pathways, consistent with potential antiviral or restorative effects, whereas natalizumab and S1P modulators showed patterns indicative of reduced immune surveillance and possible facilitation of viral persistence. Interferon isoforms and corticosteroids occupied an intermediate position, displaying mixed transcriptomic directionality depending on context and isoform.

Our transcriptome-informed classification provides a novel lens for understanding how MS therapies interact with EBV-driven disease biology. By contrasting drug-induced signatures against the convergent MS–EBV transcriptome, we identified two mechanistic categories the “reinforcers,” such as dimethyl fumarate, fingolimod, and IFN-β1b, whose transcriptomic effects aligned with EBV-associated dysregulation and the “reversers,” including cladribine, rituximab, and IFN-β1a, which shifted expression in the opposite direction. The divergence within drug classes was particularly striking, with corticosteroids and interferon isoforms splitting across both groups, underscoring that therapies considered clinically similar may exert opposite effects at the MS–EBV interface. Importantly, these classifications are consistent with established biological mechanisms. For example, rituximab and cladribine deplete EBV-harboring B cells, supporting their role as reversers, while fingolimod reduces immune surveillance, aligning with its classification as a reinforcer. Although exploratory, this framework highlights the potential of transcriptome-based classification to complement existing drug groupings and refine treatment strategies, especially in EBV-seropositive patients, by distinguishing agents that counteract EBV-driven perturbations from those that may inadvertently reinforce them.

Pathway-level analysis showed that many MS therapies reversed the MS–EBV convergent transcriptomic signature by suppressing cytokine signaling. Thirteen drugs downregulated the “cytokine–cytokine receptor interaction” pathway, while a similar set also counteracted EBV’s exploitation of this axis through cytokine mimicry by reversing the “Viral protein interaction with cytokine and cytokine receptor.” Over the course of evolution, viruses have acquired diverse strategies to evade host immunity, such as disrupting antigen presentation pathways and imitating host immune functions ^18,68^. This finding suggests that MS therapies act, at least in part, by reversing the very immune pathways hijacked by EBV to maintain persistence. In doing so, they suppress not only host-driven inflammatory overactivation but also viral evasion strategies such as cytokine mimicry, mechanisms that may contribute to EBV-driven MS pathogenesis.

Additionally, MS therapies restored pathways suppressed in the MS–EBV signature, including MAPK signaling and osteoclast differentiation. Although MAPK overactivation in microglia has been linked to neurodegeneration in MS ^69^, our network analysis, focused on the MS–EBV convergent transcriptomic signature rather than the broader MS transcriptome, revealed suppression of MAPK signaling within this overlap, with several therapies reversing this deficit. This apparent discrepancy likely arises because the KEGG MAPK pathway represents an umbrella category encompassing four distinct branches (ERK1/2, JNK, p38, and ERK5), which can be differentially regulated, overactive in microglia yet collectively appearing suppressed in the MS–EBV transcriptomic signature due to pathway-level aggregation. Moreover, evidence indicates that EBV infection can modulate MAPK regulation by downregulating ERK negative feedback phosphatases ^70^. Thus, MAPK signaling in MS is not uniformly dysregulated but instead shows branch- and cell type–specific alterations, with therapies potentially restoring EBV-suppressed modules while pathogenic overactivation persists in others.

Our analysis also revealed that a broad range of MS therapies (cladribine, dalfampridine, dimethyl fumarate, fingolimod, corticosteroids, interferons, mitoxantrone, rituximab, and teriflunomide) consistently upregulated the osteoclast differentiation pathway, which was suppressed in the MS–EBV convergent transcriptomic signature. This is notable given that MS patients are at increased risk of low bone density and altered bone homeostasis, and prior studies have shown that IFN-β treatment can modulate the RANK/RANKL/OPG axis, reducing osteoclastogenesis while promoting bone formation ^71^. Restoration of osteoclast differentiation in our analysis likely reflects normalization of pathways dysregulated by EBV and MS. Because osteoclast precursors overlap with monocyte and dendritic cell lineages, this pathway links bone metabolism with immune regulation. Its consistent reversal across therapies suggests a shared axis counteracting EBV-driven immune disruption, though clinically some agents, notably corticosteroids, are also associated with osteoporosis. This paradox highlights the dual role of osteoclast pathways in immune defense and bone turnover, where therapeutic modulation may both restore antiviral signaling and increase skeletal risk.

Notably, IL-17, TNF, and Toll-like receptor signaling appeared in both categories of pathway reversal, with the same therapies associated with downregulation of genes pathologically upregulated in the MS–EBV signature and simultaneous upregulation of genes that were abnormally suppressed. This bidirectional regulation suggests that MS immunotherapies do not simply switch entire pathways on or off, but rather recalibrate them by suppressing hyperinflammatory modules while reactivating components important for antiviral defense and tissue homeostasis.

Finally, our genotype-informed network pharmacology framework revealed how MS therapies intersect with the viral–host interface at genetic susceptibility nodes. Dimethyl fumarate and natalizumab showed the broadest coverage and S1P modulators the least. Strikingly, many therapies converged on *HLA-DRB1*, the strongest genetic risk factor for MS, which not only presents EBV-derived peptides ^11,12^ but also functions as an EBV entry cofactor ^13^. This emphasizes its role as a shared node across T cell–targeting agents, glatiramer acetate, dimethyl fumarate, natalizumab, and anti-CD20 antibodies. Beyond HLA-DRB1, variants in *IL7R, TYK2, STAT3, CASP8, CD40*, and *TNFRSF1A* were frequently engaged, highlighting convergence on immune regulators central to both MS susceptibility and EBV biology. Several drugs, including dimethyl fumarate, natalizumab, alemtuzumab, and glatiramer acetate, targeted pathways that link MS risk variants directly to EBV processes, such as phagosome-mediated antigen presentation, NF-κB signaling, and lipid metabolism, pathways exploited by EBV during latency and reactivation. These findings suggest that MS therapies not only modulate immunity broadly but also intersect with host–virus interactions at genetically defined nodes. By repeatedly converging on susceptibility alleles like HLA-DRB1, immunotherapies reinforce a genetic–viral axis in MS pathogenesis, providing a framework to interpret therapeutic impact in relation to both host genetics and EBV manipulation. Collectively, these observations illustrate how a virus-informed pharmacogenomic framework can uncover therapeutic mechanisms that remain hidden in conventional immunological or genetic analyses.

Our analysis is not without limitations. Pathway enrichment was used to infer drug effects from transcriptomic signatures, but this limits interpretability. Specifically, including all differentially expressed genes obscures directionality, while separating up- and downregulated sets can yield contradictions, as seen with IL-17, TNF, and Toll-like receptor signaling, indicating inherent limitations of current enrichment methods. Second, drug–target associations were derived from KEGG and DrugBank, which capture only established interactions; additional context-specific or unknown off-target effects are not represented. Third, the reinforcer/reverser assignments did not survive multiple-testing correction, and some pathways appeared regulated in both directions, underscoring the complexity of capturing system-level drug effects. Fourth, MS risk genes were obtained from DisGeNET, which may not fully represent all known loci and exclude undiscovered susceptibility variants. Finally, the framework infers EBV-linked interactions but lacks patient-level measures such as viral load, genetics, immune profiles, imaging, or clinical outcomes; integration of such multi-omic datasets would enable validation but remains limited by scarce EBV-specific data and fragmented cohorts, as is also the case for most clinical and preclinical studies in this field.

Despite these limitations, our virus-informed network pharmacology framework provides an important proof-of-principle for mapping how existing MS immunotherapies intersect with EBV-driven mechanisms. By embedding genetic susceptibility, viral protein interactions, and drug-induced transcriptomes within a single framework, we highlight points of convergence that are not accessible through traditional models. Therapies with broad network engagement (e.g., dimethyl fumarate, natalizumab, interferons) may most profoundly influence EBV-driven MS biology, for better or worse. Narrower agents (e.g., anti-CD20 antibodies, S1P modulators) act more selectively, often via reservoir depletion or trafficking control, but may fail to rewire EBV–host interactions at the systems level. Finally, the reinforcer versus reverser classification provides a novel lens to distinguish therapies not only by immunological class but also by their alignment with or opposition to EBV-driven immune signatures. By linking viral, genetic, and pharmacologic dimensions, this framework suggests a path toward precision medicine in MS, where therapeutic selection could one day be tailored to an individual’s EBV status, genetic risk, and immune network profile.

## 5. Conclusion

This study shows that MS immunotherapies vary widely in their engagement with the MS–EBV network, genetic risk variants, and viral processes. Dimethyl fumarate, natalizumab, and interferons exert the broadest effects, while S1P modulators and anti-CD20 antibodies act more narrowly. Importantly, multiple therapies intersected with HLA-DRB1, NF-κB, and cytokine signaling, underscoring shared points of convergence between host genetics, viral manipulation, and therapeutic action. At the transcriptomic level, immunotherapies stratified into “reinforcers” and “reversers” of the convergent MS–EBV signature, revealing that even drugs within the same clinical class can exert divergent transcriptomic influences. Across classes, however, a consistent reversal of EBV-exploited cytokine signaling pathways was observed, suggesting a unifying therapeutic mechanism that mitigates viral persistence and immune dysregulation. This study moves beyond recognizing the immunosuppression paradox to mechanistically dissect how distinct MS immunotherapies interface with EBV-driven pathways. It demonstrates that the paradox arises from drug-specific modulation of viral and host signaling networks, some restoring antiviral defense while others reinforce EBV-associated dysregulation. These insights provide a mechanistic blueprint revealing the specific vulnerabilities and strengths of each MS immunotherapy in relation to EBV-driven mechanisms, highlighting which therapeutic actions must be complemented to simultaneously suppress inflammation and control viral persistence, thereby defining a path toward EBV-informed precision immunotherapy in MS. Moreover, these insights support the rationale for developing adjunct strategies that combine immune-modulating and EBV-targeted interventions to achieve more comprehensive control of inflammation, viral persistence, and relapse prevention in MS.

## 6. Funding

This publication was made possible by support from the IDSA Foundation. Its contents are solely the responsibility of the authors and do not necessarily represent the official views of the IDSA Foundation.

## 7. Competing Interests

The author declares no competing interests. The code developed for this study is the intellectual property of the author and has not been commercially licensed or shared.

## 8. Data Availability Statement

All data used in this study are publicly available from the resources and references cited within the Methods section.

## 9. Code Availability Statement

The code used in this study was developed independently by the author and constitutes the author’s intellectual property. While the method is fully described in the manuscript, the original code is not publicly available but may be shared upon reasonable request, subject to intellectual property considerations.

## Notes

### Competing Interest Statement

The authors have declared no competing interest.

